# Efficient cloning of genes for wheat yield component traits from QTLs via sequencing of the RIL population

**DOI:** 10.1101/2024.02.22.574000

**Authors:** Mingxia Zhang, Xu Han, Hui Wang, Junsheng Sun, Baojin Guo, Minggang Gao, Huiyan Xu, Guizhi Zhang, Hongna Li, Xiaofeng Cao, Nannan Li, Yiru Xu, Qun Wu, Chunyang Wang, Guohua Zhang, Yapei Yuan, Junxia Man, Yanyan Pu, Guangde Lv, Chunyan Qu, Jinjie Sun, Xiyong Cheng, Xinjuan Dong, Fanmei Kong, Yan Zhao, Yanrong An, Yuanyuan Yuan, Ying Guo, Sishen Li

## Abstract

In wheat (*Triticum aestivum* L.), yield component traits (YCTs) are the most important yield traits. Only several genes for YCTs have been originally cloned. The efficient cloning of genes for YCTs directly from wheat remains a challenge. Here, we proposed a strategy for cloning genes from quantitative trait loci (QTLs) by sequencing of recombinant inbred lines (RILs) (QTL-Seq-RIL). Using the ‘TN18 × LM6’ RIL population as an example, we identified 138 candidate unigenes (CUGs) for YCTs from 77 stable QTLs. The average of CUGs per QTL was 1.8, which enabled us to confirm the CUGs directly. We have confirmed seven CUGs, *TaIFABPL, TaDdRp, TaRLK, TaTD, TaTFC3, TaKMT* and *TaSPL17*, via the CRISPR/Cas9 system. Of these, six genes were found firstly to regulate YCTs in crops except for *TaSPL17*. Five CUGs (include *TaSPL17*) for which orthologous genes have been cloned previously with the same or similar agronomic functions. It is to say, 11 CUGs were preliminarily validated using a single RIL population. QTL-Seq-RIL provides an efficient method for rapid gene cloning using existing RIL populations.

## Introduction

Common wheat (*Triticum aestivum* L., 2n=6x=42, AABBDD) is one of the most important cereal crops worldwide, providing approximately one-fifth of all calories consumed by humans (FAO, http://www.fao.org/faostat/). The demand for wheat is projected to increase 60% by 2050 to feed a population of 9-10 billion people (Godfray et al., 2010). Increasing grain yield is therefore an urgent task to ensure global food and nutritional security. The most important yield traits are yield component traits (YCTs), including spike number per unit area (SN), grain number per spike (GNS) and thousand-grain weight (TGW). Cloning the genes regulating YCTs is a key question for wheat genetic improvement.

Obtaining a sufficient number of genes is a pre-requisite for further research on action mechanisms and variety improvement. In wheat, the less number of cloned genes is a bottleneck for gene-related research. The wheat genes originally cloned were mainly obtained by the traditional forward genetic strategy of ‘map-based cloning’, such as *VRN* (Yan et al., 2003), *Ms2* (Ni et al., 2017), *Gpc-B1* (Uauy et al., 2006) and some disease resistance genes (Feuillet et al., 2003; Fu et al., 2009; Periyannan et al., 2013; Li et al., 2019; Su et al., 2019; Lu et al., 2020). Map-based cloning, however, is laborious and time-consuming and has been limited in wheat due to its allohexaploid nature with a large (∼17 Gb) and highly repetitive genome (IWGSC, 2014).

Based on genome sequencing, some rapid wheat disease resistance gene cloning strategies have been proposed recently. Sanchez-Martin et al. (2016) developed mutant chromosome sequencing (MutChromSeq) to identify induced mutations, and applied it to clone the barley *Eceriferum-q* and the wheat *Pm2* genes. Steuernagel et al. (2016) described a three-step method MutRenSeq that combines chemical mutagenesis with exome capture and sequencing for rapid disease resistance gene cloning, and applied it to clone stem rust resistance genes *Sr22* and *Sr45* from wheat. Thind et al. (2017) reported targeted chromosome-based cloning via long-range assembly (TACCA), and cloned the *Lr22a* leaf-rust resistance gene. Arora et al. (2019) combined association genetics with R gene enrichment sequencing (AgRenSeq) to exploit pan-genome variation in wild diploid wheat, and rapidly cloned four stem rust resistance genes. Athiyannan et al. (2022) generated a chromosome-scale assembly of a wheat cultivar by combining high-fidelity long reads, optical mapping and chromosome conformation capture, and identified the *Yr27* gene.

YCTs are the most complex quantitative traits controlled by multiple genes and are highly influenced by environmental conditions. As far as we know, only eight genes for YCTs have been originally cloned in wheat, which the functions were verified using transgenic technologies, via map-based cloning, bulked segregant analysis (BSA) and genome-wide association study (GWAS). The efficient cloning of genes for YCTs directly from wheat remains a challenge. For GNS related traits, four genes were originally cloned using map-based cloning strategy. *GNI1* gene, encoding an HD-Zip I transcription factor, was identified as responsible for grain number per spikelet. The function of *GNI1* was verified by the RNAi-based knockdown (Sakuma et al., 2019). *WFZP* is a causal gene for triple spikelet. CRISPR/Cas9-based gene mutation indicated that *WFZP* simultaneously controls spikelet formation and awn elongation (Du et al., 2021). *FT-D1* is a well-known flowering time gene in wheat. Analysis of *FT-D1* mutant lines developed by CRISPR/Cas9 provided genetic evidence for the function in regulating spikelet number and heading date (Chen et al., 2022). *TaCOL-B5*, encoding a CONSTANS-like (COL) protein, is a gene determining the number of spikelet nodes per spike in common wheat, which was verified via overexpression and gene editing (Zhang et al., 2022).

For TGW related traits, three genes were originally cloned. *Tasg-D1*, the grain-shape gene in *T. sphaerococcum*, was isolated via map-based cloning. *Tasg-D1* encodes a Ser/Thr protein kinase glycogen synthase kinase3 (STKc_GSK3). The function of *Tasg-D1* was confirmed by ethyl methanesulfonate (EMS) mutagenesis and overexpression (Cheng et al., 2020). *KAT-2B* was cloned through a *tgw1* mutant by bulked segregant analysis (BSA). The biological function of *KAT-2B* in controlling TGW was confirmed by overexpression in wild type, *tgw1*, and *Arabidopsis kat2* mutant (Chen et al., 2020). *ALI-1*, which regulates awn elongation and TGW, was identified by genome-wide association study (GWAS) and fine mapping. *ALI-1* was confirmed to be the underlying gene of the *B1* locus through the functional complimentary test with native awnless allele (Wang et al., 2020).

For SN related traits, only one gene *TN1* was originally cloned using map-based cloning. *TN1* encodes a transmembrane ankyrin repeat protein. A genetic complementation test and CRISPR/Cas9-mediated genome editing confirmed *TN1* gene is responsible for the low-tillering phenotype of the *tn1* mutant. (Dong et al., 2023).

Here, combining forward genetics with genome re-sequencing technology, we propose a strategy for cloning genes from quantitative trait loci (**<u>QTL</u>**s) by **<u>seq</u>**uencing of the recombinant inbred line (**<u>RIL</u>**) population (QTL-Seq-RIL). This approach involves sequencing the parents and each line of the RILs via RNA-Seq, and sequencing parents and a subset of RILs via DNA-Seq. Using the ‘TN18 × LM6’ RIL population (TL-RIL) as an example, we demonstrate the detailed results and efficacy of the QTL-Seq-RIL, showcasing its potential in the field of gene cloning.

## Results

### Overall procedure for gene cloning strategy of QTL-Seq-RIL

We proposed a QTL-Seq-RIL strategy of clone genes from RIL popuatiom. The overall procedure is: 1) profiling each RIL for RNA-Seq; 2) assembling reads according to the reference genome, and calling SNPs/InDels of each RIL; 3) merging SNPs/InDels into sub-unigenes and constructing a genetic map of unigenes (UG-Map) based on the physical position of unigenes; 4) mapping QTLs based on the UG-Map and phenotypic data in order to identify stable QTLs; 5) finding the candidate unigenes (CUGs) within the interval of corresponding QTLs, and identifying the mutants in promoters of CUGs by DNA sequencing of parents and a subset of RILs; and 6) confirming CUGs using gene editing system known as CRISPR/Cas9, etc. (Fig. 1A; also reference to Materials and Methods section). Using QTL-Seq-RIL, many polymorphic CUGs of stable QTLs for target traits in a RIL popultion can be identified. Then, some CUGs can be selected to confirm the gene functions (Fig. 1B).

**Fig. 1.**
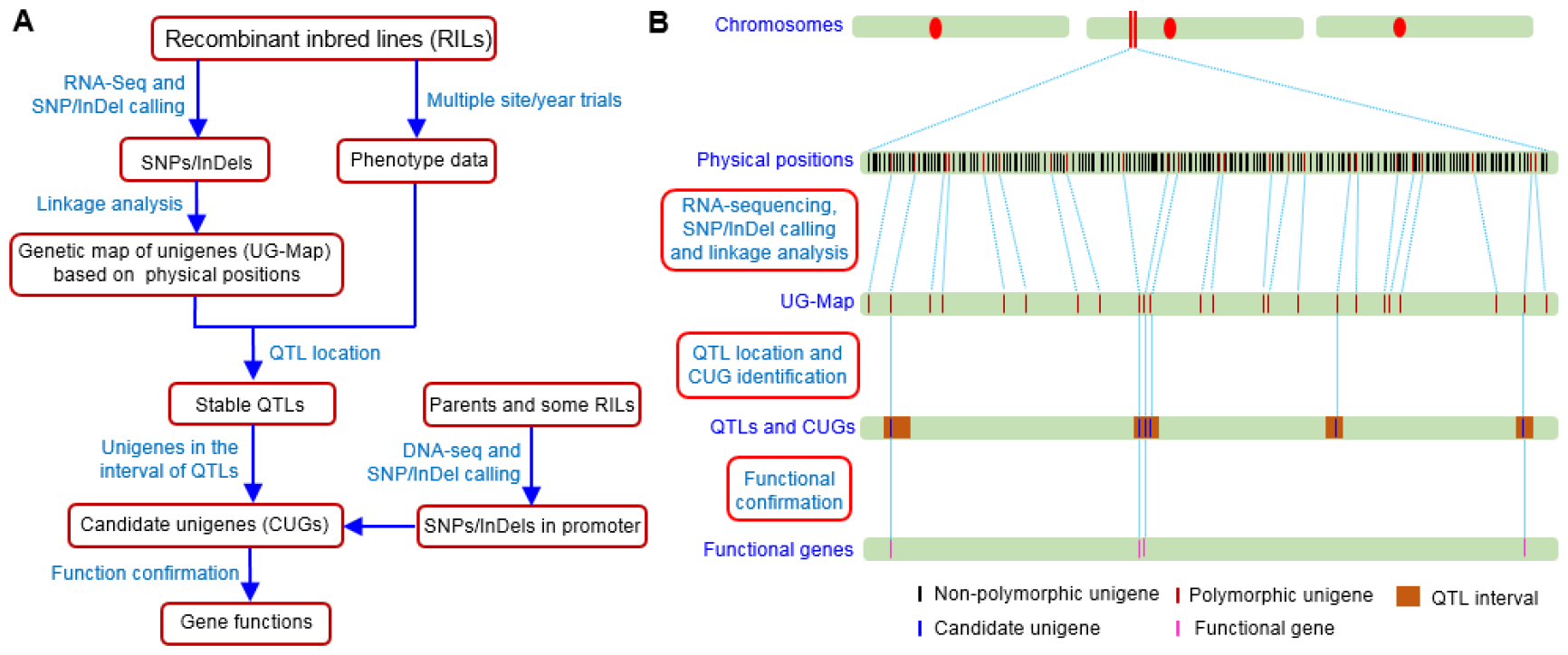
Strategy for cloning genes from QTLs by sequencing of the RILs (QTL-Seq-RIL). **A**, Steps of the QTL-RNASeq-RIL strategy. **B**, Diagrammatic sketch from all unigenes in genome to functional genes.

Using the TL-RILs as an example, we constructed a UG-Map with 3,1445 sub-unigenes, identified 138 CUGs for YCTs from 77 stable QTLs, and confirmed the functions of seven CUGs using the CRISPR/Cas9 system. The detailed results are as follows.

### RNA sequencing and UG-Map construction

Clean data were obtained for TN18, LM6 and each line of TL-RILs using RNA-Seq technology. By mapping to the RefSeq v1.1 reference genome (IWGSC, 2018), we found 4.10E+08, 1.48E+08 and 1.08E+10 unique reads for TN18, LM6 and TL-RILs, respectively. After SNP/InDel calling, genotype merging and filtering, a total of 31,445 polymorphic sub-unigenes were obtained. The sub-unigenes showed high levels of recombination among the RILs, with several recombination hotspots on each chromosome, distributed mainly at the ends or near the ends of chromosomes (Fig. S1).

The polymorphic sub-unigenes were used to construct the UG-Map, and the final UG-Map included 31,445 sub-unigenes, which represented 27,452 sites, 27,983 unigenes and 118,120 SNPs/InDels (Fig. 2A, 2B; Table 1, S1). Among the unigenes, 21,063 were annotated in RefSeq v1.1 (including 16,267 high confidence (HC) genes and 4,796 low confidence (LC) genes), 2,545 were newly annotated in the TL-RILs (named as *Traes****TL****1A02G00100*, etc.), and the other 4,375 were non-coding RNAs (ncRNAs, named as *STRG_1A*.*000001*, etc). The number of polymorphic unigenes per chromosome ranged form 387 of 3D to 2,581of 1B; and 14,328, 9,612 and 4,043 unigenes were on sub-genomes B, A and D, respectively.

**Table 1.** Summary of the UG-Map for the TL-RILs. The UG-Map was constructed using sub-unigenes.

| Chromosome | Number of sites | Number of SNPs/InDels | Number of unigenes/sub-unigenes |  |  |  |  |  |  |  |
| --- | --- | --- | --- | --- | --- | --- | --- | --- | --- | --- |
|  |  |  | Total |  | RefSeq v1.1 |  | TL-RIL |  | ncRNA |  |
|  |  |  | unigene | sub-unigene | unigene | sub-unigene | unigene | sub-unigene | unigene | sub-unigene |
| 1A | 1306 | 5346 | 1277 | 1470 | 966 | 1114 | 119 | 129 | 192 | 227 |
| 1B | 2379 | 11673 | 2581 | 2824 | 1928 | 2072 | 245 | 273 | 408 | 479 |
| 1D | 875 | 4190 | 897 | 976 | 704 | 759 | 87 | 91 | 106 | 126 |
| 2A | 2047 | 8565 | 2068 | 2389 | 1459 | 1719 | 213 | 229 | 396 | 441 |
| 2B | 1674 | 6652 | 1597 | 1848 | 1221 | 1399 | 148 | 171 | 228 | 278 |
| 2D | 984 | 4706 | 941 | 1083 | 769 | 878 | 75 | 85 | 97 | 120 |
| 3A | 890 | 2930 | 808 | 966 | 588 | 683 | 68 | 75 | 152 | 208 |
| 3B | 2049 | 8969 | 2091 | 2346 | 1548 | 1702 | 187 | 206 | 356 | 438 |
| 3D | 447 | 1071 | 387 | 462 | 298 | 363 | 38 | 41 | 51 | 58 |
| 4A | 1179 | 4079 | 1118 | 1312 | 841 | 957 | 110 | 128 | 167 | 227 |
| 4B | 1295 | 6373 | 1461 | 1565 | 1139 | 1212 | 120 | 128 | 202 | 225 |
| 4D | 268 | 630 | 267 | 301 | 202 | 223 | 15 | 15 | 50 | 63 |
| 5A | 1526 | 7469 | 1594 | 1750 | 1246 | 1363 | 123 | 133 | 225 | 254 |
| 5B | 2333 | 12003 | 2579 | 2778 | 1958 | 2097 | 233 | 244 | 388 | 437 |
| 5D | 936 | 2926 | 939 | 1051 | 738 | 820 | 68 | 72 | 133 | 159 |
| 6A | 1491 | 8041 | 1605 | 1756 | 1202 | 1315 | 147 | 155 | 256 | 286 |
| 6B | 1776 | 7362 | 1877 | 2082 | 1367 | 1495 | 184 | 207 | 326 | 380 |
| 6D | 633 | 1540 | 612 | 707 | 445 | 512 | 52 | 59 | 115 | 136 |
| 7A | 1226 | 5319 | 1142 | 1330 | 849 | 992 | 112 | 120 | 181 | 218 |
| 7B | 1053 | 4536 | 997 | 1154 | 714 | 802 | 113 | 122 | 170 | 230 |
| 7D | 1085 | 3740 | 1145 | 1295 | 881 | 986 | 88 | 96 | 176 | 213 |
| Sub-genome A | 9665 | 41749 | 9612 | 10973 | 7151 | 8143 | 892 | 969 | 1569 | 1861 |
| Sub-genome B | 13644 | 61308 | 14328 | 15892 | 10756 | 11765 | 1318 | 1447 | 2254 | 2680 |
| Sub-genome D | 4143 | 15063 | 4043 | 4580 | 3156 | 3555 | 335 | 363 | 552 | 662 |
| Total | 27452 | 118120 | 27983 | 31445 | 21063 | 23463 | 2545 | 2779 | 4375 | 5203 |

**Fig. 2.**
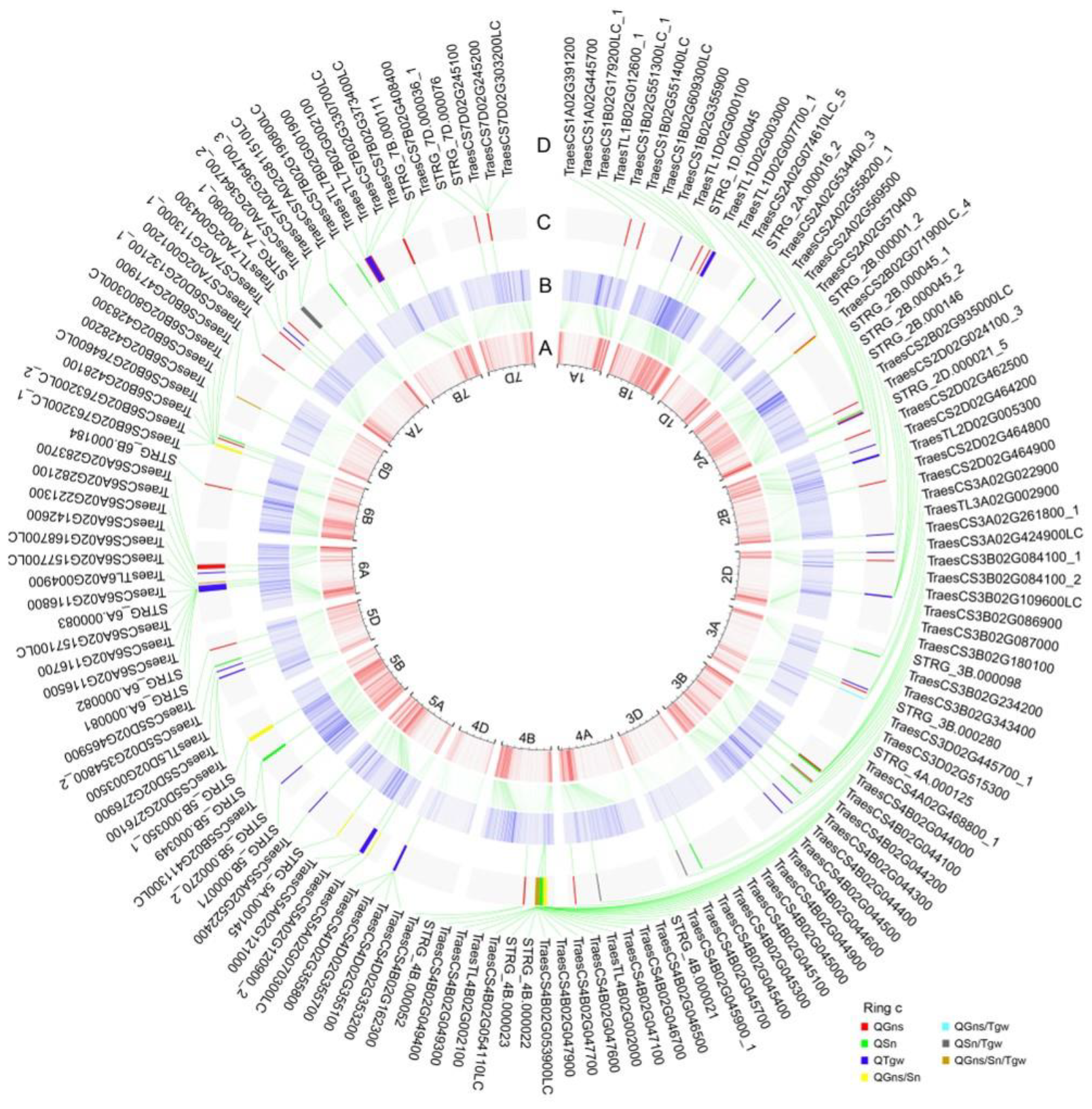
Physical positions of the sub-unigenes, UG-Map, QTLs and their CUGs of TL-RILs. **A**, Physical position of the 31,445 sub-unigenes in RefSeq v1.1 with one scale representing 100 Mb. **B**, Genetic position (cM) of the 31,445 sub-unigenes in the UG-Map. **C**, Distribution of the 77 QTLs on the 21 chromosomes for yield component traits. **D**, Distribution of the 196 sub-unigenes for corresponding QTLs. The rings between **A** and **B, B** and **C**, and **C** and **D** are the linking lines of the corresponding positions.

### QTL detection and CUG identification

The phenotypic data of the TL-RILs and their parents for SN, GNS and TGW were investigated under 22 environments/treatments (Table S2). Combining the UG-Map and phenotypic data, a total of 77 stable QTLs for the three yield traits were detected on all 21 chromosomes. (Fig. 2C; Table S3). Forty-eight QTLs for 51 traits (some QTLs related more than one traits) showed positive additive effects, with the female parent TN18 increasing the QTL effects; whereas 40 QTLs for 41 traits showed negative additive effects, with the male parent LM6 increasing the QTL effects. A number of 64 QTLs were associated solely with one trait (Table 2, S3). Of these, 27 QTLs for GNS were located on 15 chromsomes, 1A, 1B, 2A, 2B, 2D, 3A, 3B, 4A, 4B, 5D, 6A, 6B, 7A, 7B and 7D. For SN, 13 QTLs were obtained on 11 chromsomes, 1D, 2A, 3A, 3B, 3D, 4B, 5B, 5D, 6B, 7A and 7B. For TGW, 24 QTLs were found on 14 chromsomes, 1B, 1D, 2A, 2B, 2D, 3A, 3B, 4D, 5A, 5B, 5D, 6A, 7A and 7B. The other 13 QTLs were associated with two or three traits, including 3, 6, 1 and 3 QTL(s) for GNS/SN/TGW, GNS/SN, GNS/TGW and SN/TGW, respectively.

**Table 2.** Summary of 77 stable QTLs and their 138 CUGs. *TraesTL*, unigenes annotated in the TL-RILs; *STRG*, ncRNA.

| QTL | Unigene | QTL | Unigene | QTL | Unigene | QTL | Unigene |
| --- | --- | --- | --- | --- | --- | --- | --- |
| *QGns-1A-10882* | *TraesCS1A02G391200* | *QSn-3B-4564* | *TraesCS3B02G109600LC* |  | *STRG_4B.000022* |  | *TraesCS6A02G157700LC* |
| *QGns-1A-13226* | *TraesCS1A02G445700* |  | *TraesCS3B02G086900* |  | *STRG_4B.000023* | *QTgw-6A-5080* | *TraesCS6A02G168700LC* |
| *QTgw-1B-4752* | *TraesCS1B02G179200LC* |  | *TraesCS3B02G087000* |  | *TraesCS4B02G054110LC* | *QGns/Sn/Tgw-6A-5634* | *TraesCS6A02G142600* |
| *QGns-1B-9098* | *TraesTL1B02G012600* | *QGns-3B-6834* | *TraesCS3B02G180100* |  | *TraesTL4B02G002100* | *QTgw-6A-7089* | *TraesCS6A02G221300* |
| *QGns-1B-10676* | *TraesCS1B02G551300LC* | *QSn-3B-7126* | *STRG_3B.000098* |  | *TraesCS4B02G049300* | *QGns-6A-8001* | *TraesCS6A02G282100* |
|  | *TraesCS1B02G551400LC* | *QSn-3B-9800* | *TraesCS3B02G234200* |  | *TraesCS4B02G049400* |  | *TraesCS6A02G283700* |
| *QTgw-1B-11562* | *TraesCS1B02G609300LC* | *QTgw-3B-12014* | *TraesCS3B02G343400* | *QGns-4B-3410* | *STRG_4B.000052* | *QGns-6B-7868* | *STRG_6B.000184* |
|  | *TraesCS1B02G355900* | *QTgw-3B-14462* | *STRG_3B.000280* |  | *TraesCS4B02G162300* | *QGns/Sn-6B-15200* | *TraesCS6B02G763200LC* |
| *QSn-1D-508* | *TraesTL1D02G000100* | *QSn-3D-4808* | *TraesCS3D02G445700* | *QTgw-4D-3090* | *TraesCS4D02G353200* |  | *TraesCS6B02G428100* |
| *QTgw-1D-4402* | *STRG_1D.000045* | *QSn/Tgw-3D-6218* | *TraesCS3D02G515300* |  | *TraesCS4D02G355100* |  | *TraesCS6B02G764600LC* |
|  | *TraesTL1D02G003000* | *QSn/Tgw-4A-9135* | *STRG_4A.000125* |  | *TraesCS4D02G355700* |  | *TraesCS6B02G428200* |
| *QTgw-1D-6724* | *TraesTL1D02G007700* | *QGns-4A-12540* | *TraesCS4A02G468800* |  | *TraesCS4D02G355800* |  | *TraesCS6B02G428300* |
| *QGns-2A-1604* | *TraesCS2A02G074610LC* | *QGns/Sn-4B-1132* | *TraesCS4B02G044000* | *QGns/Sn-5A-1720* | *TraesCS5A02G070300LC* | *QGns-6B-16461* | *TraesCS6B02G800300LC* |
| *QGns/Sn-2A-1671* | *STRG_2A.000016* |  | *TraesCS4B02G044100* | *QTgw-5A-2613* | *TraesCS5A02G120900* | *QSn-6B-16986* | *TraesCS6B02G471900* |
| *QGns-2A-16450* | *TraesCS2A02G534400* |  | *TraesCS4B02G044200* |  | *TraesCS5A02G121000* | *QGns/Sn/Tgw-6D-3306* | *TraesCS6D02G132100* |
| *QSn-2A-18173* | *TraesCS2A02G558200* |  | *TraesCS4B02G044300* | *QGns/Sn-5A-6640* | *STRG_5A.000145* | *QGns-7A-0* | *TraesCS7A02G001200* |
| *QGns-2A-18533* | *TraesCS2A02G569500* |  | *TraesCS4B02G044400* | *QTgw-5A-11665* | *TraesCS5A02G522400* | *QGns-7A-4157* | *TraesCS7A02G113000* |
| *QTgw-2A-18633* | *TraesCS2A02G570400* |  | *TraesCS4B02G044500* | *QTgw-5B-2761* | *STRG_5B.000071* | *QTgw-7A-5018* | *TraesTL7A02G004300* |
| *QGns-2B-48* | *STRG_2B.000001* |  | *TraesCS4B02G044600* | *QSn-5B-7525* | *STRG_5B.000270* | *QGns-7A-6117* | *STRG_7A.000080* |
| *QTgw-2B-2772* | *TraesCS2B02G071900LC* | *QSn-4B-1170* | *TraesCS4B02G044900* |  | *TraesCS5B02G411300LC* | *QSn/Tgw-7A-9048* | *TraesCS7A02G364700* |
| *QTgw-2B-4768* | *STRG_2B.000045* |  | *TraesCS4B02G045000* | *QGns/Sn-5B-11490* | *STRG_5B.000349* | *QSn-7A-14459* | *TraesCS7A02G811510LC* |
| *QGns-2B-9522* | *STRG_2B.000146* |  | *TraesCS4B02G045100* |  | *STRG_5B.000350* | *QSn-7B-2205* | *TraesCS7B02G190800LC* |
| *QTgw-2B-19162* | *TraesCS2B02G935000LC* |  | *TraesCS4B02G045300* | *QTgw-5D-4112* | *TraesCS5D02G276100* | *QGns-7B-3280* | *TraesTL7B02G001900* |
| *QTgw-2D-576* | *TraesCS2D02G024100* |  | *TraesCS4B02G045400* |  | *TraesCS5D02G276900* | *QTgw-7B-3580* | *TraesTL7B02G002100* |
| *QGns-2D-1629* | *STRG_2D.000021* |  | *TraesCS4B02G045700* | *QTgw-5D-5021* | *TraesTL5D02G003500* |  | *TraesCS7B02G330700LC* |
| *QTgw-2D-6108* | *TraesCS2D02G462500* |  | *TraesCS4B02G045900* | *QSn-5D-5788* | *TraesCS5D02G354800* | *QGns-7B-3744* | *TraesCS7B02G373400LC* |
|  | *TraesTL2D02G005300* | *QGns/Sn/Tgw-4B-1220* | *STRG_4B.000021* | *QGns-5D-8048* | *TraesCS5D02G465900* | *QGns-7B-8140* | *STRG_7B.000111* |
|  | *TraesCS2D02G464200* |  | *TraesCS4B02G046500* | *QTgw-6A-4765* | *STRG_6A.000081* |  | *TraesCS7B02G408400* |
|  | *TraesCS2D02G464800* |  | *TraesCS4B02G046700* |  | *STRG_6A.000082* | *QGns-7D-4272* | *STRG_7D.000036* |
|  | *TraesCS2D02G464900* |  | *TraesCS4B02G047100* |  | *TraesCS6A02G157100LC* | *QGns-7D-6025* | *STRG_7D.000076* |
| *QSn-3A-1163* | *TraesCS3A02G022900* |  | *TraesTL4B02G002000* |  | *TraesCS6A02G116500* |  | *TraesCS7D02G245100* |
| *QTgw-3A-5250* | *TraesTL3A02G002900* |  | *TraesCS4B02G047600* |  | *TraesCS6A02G116700* |  | *TraesCS7D02G245200* |
| *QGns-3A-5650* | *TraesCS3A02G261800* |  | *TraesCS4B02G047700* |  | *STRG_6A.000083* |  | *TraesCS7D02G303200LC* |
| *QGns/Tgw-3A-6404* | *TraesCS3A02G424900LC* |  | *TraesCS4B02G047900* |  | *TraesCS6A02G116800* |  |  |
| *QGns-3B-4420* | *TraesCS3B02G084100* |  | *TraesCS4B02G053900LC* |  | *TraesTL6A02G004900* |  |  |

A total of 138 CUGs (142 sub-unigenes) were identified from 77 stable QTLs. Of these, 34, 22, 44, 17, 17, 1 and 3 CUG(s) were related to GNS, SN, TGW, GNS/SN/TGW, GNS/SN, GNS/TGW and SN/TGW, respectively (Fig. 2D; Table 2, S3). Among the 138 CUGs, 99 were annotated in RefSeq v1.1 (including 75 HC and 24 LC genes), 13 were annotated in TL-RILs, and the other 26 were ncRNAs. The average number of CUGs per QTL was 1.8, with 57 QTLs containing only one CUG, 11 QTLs containing two CUGs, and the other eight QTLs containing 3-15 CUGs (Table 2, S3).

For the 99 CUGs annotated in RefSeq v1.1, 50 and 20 were non-synonymous and synonymous mutations in exons; and 14 and 15 were mutant in UTRs and in introns, respectively. For all 138 CUGs, the reads of 97 CUGs were significantly different between the TN18 and LM6 types in TL-RILs, indicating that their mRNA expression levels were changed (Table S4). According to the DNA sequences of the parents and 18 RILs, the promoter region (−2000 bp from the start site of the 5’UTR) of 42 of the 99 CUGs in RefSeq v1.1 and seven of the 13 CUGs in TL-RILs were mutated (Table S5).

### *TaIFABPL* gene for *QTgw-3B-12014* regulated TGW

The meta-QTL interval of *QTgw-3B-12014* for TGW was at the 12,007.0-12,017.4 cM on chromosome 3B in the UG-Map. This region included only one CUG, *TraesCS3B02G343400* (Fig. 3A; Table 2, S3). *TraesCS3B02G343400* is annotated as an IFA (intermediate filament antigen)-binding protein-like (*TaIFABPL-B1* gene) with one InDel in 3’UTR and no SNP/InDel in promoter. The expression levels were significantly different between the TN18 and LM6 genotypes of the TL-RILs (Table S4, S5). We performed gene editing of *TaIFABPL-B1*. Three homozygous mutant genotypes, *bb-1* (subgenome B, -4 bp), *bb-2* (−15 bp) and *bb-3* (−18 bp), were identified in T_4_ generation (Fig 4A, S2A). The TGW values of *bb-1, bb-2, bb-3* and WT were 43.96, 41.99, 43.15 and 38.49 g, respectively. The TGW of mutants *bb-1, bb-2* and *bb-3* were significantly higher than that of WT by 5.47 (14.22%), 3.50 (9.10%) and 4.66 g (12.10%), respectively (Fig. 4A; Table S6). The results indicated that *TaIFABPL-B1* gene regulated TGW.

**Fig. 3.**
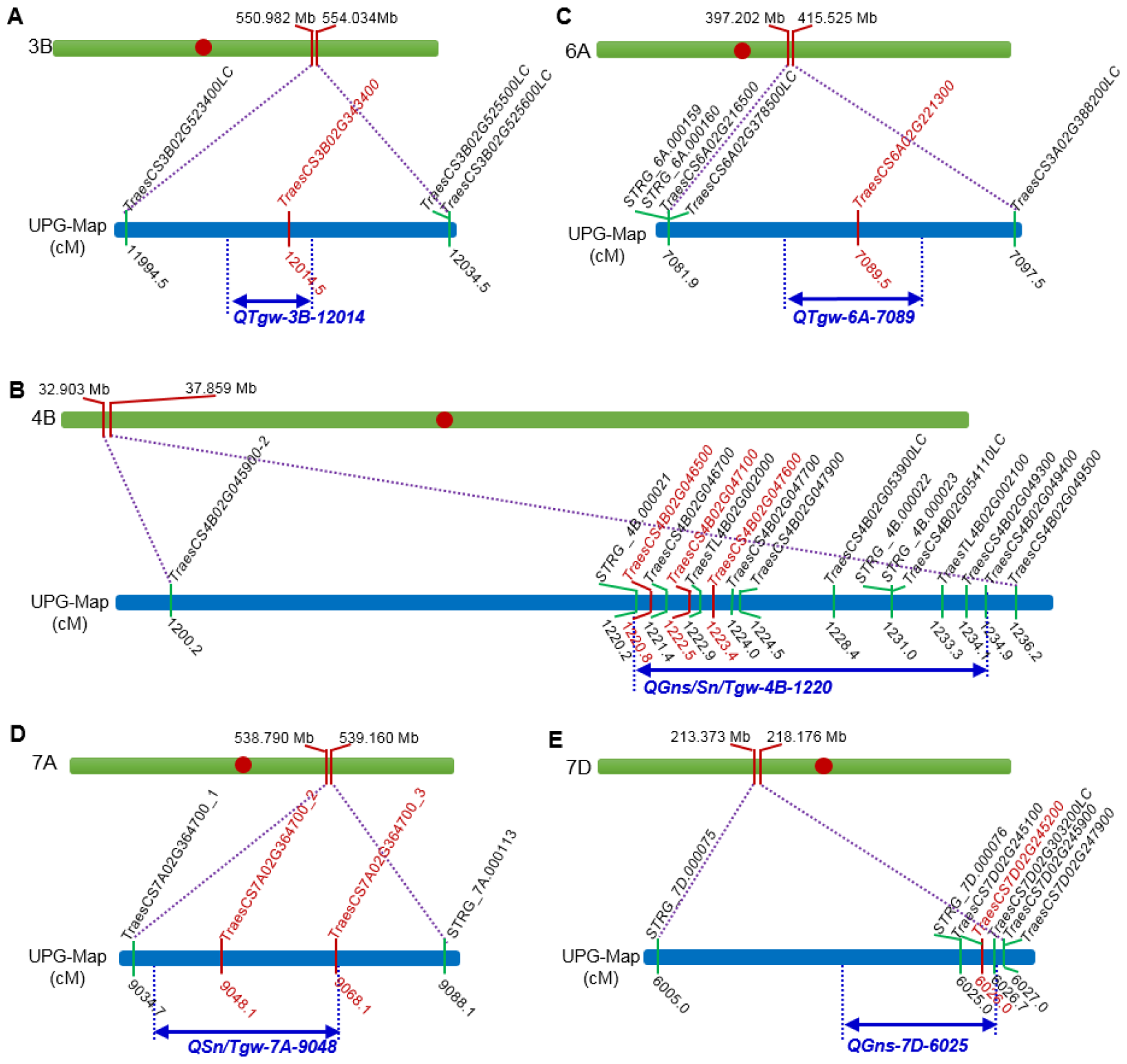
Location of the four QTLs on the UPG-Map. For each QTL, the figure included physical region of the QTL on the chromosome, genetic region of the QTL in the UPG-Map. The red CUGs were edited using CRISPR/Cas9 system. **A**, Physical and genetic region of *QTgw-3B-12014*, which included only one CUG. **B**, Physical and genetic region of *QGns/Sn/Tgw-4B-1220*, which included 14 CUGs. **C**, Physical and genetic region of *QTgw-6A-7089*, which included only one CUG. **D**, Physical and genetic region of *QSn/Tgw-7A-9048*, which included only one CUG. **E**, Physical and genetic region of *QGns-7D-6025*, which included three CUGs.

**Fig. 4.**
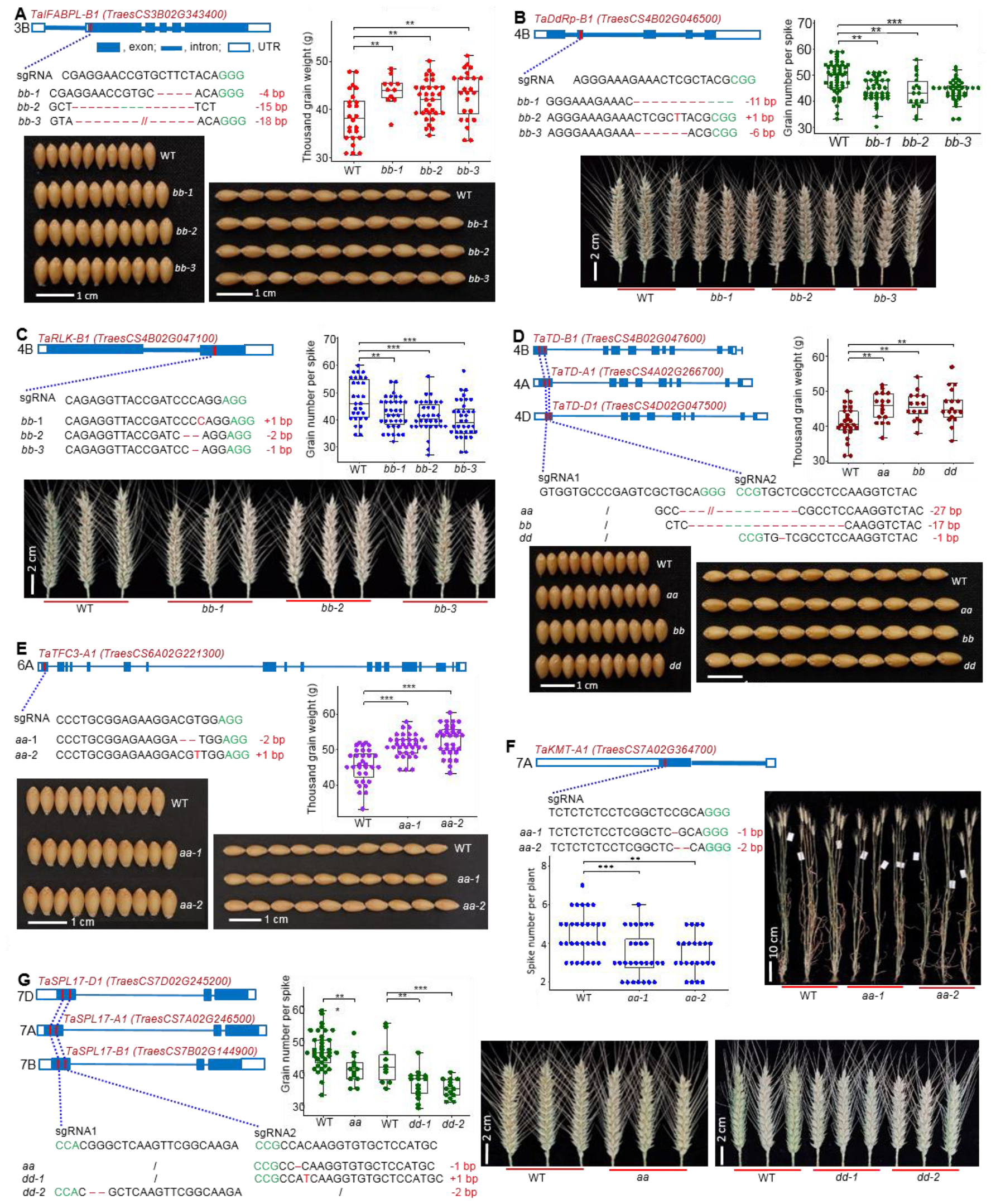
Functional confirmation of the five CUGs using CRISPR/Cas9 system. For each gene, the figure included gene structure framework, sgRNA, mutant genotypes, pictures and box plot for corresponding trait. **A**, *TaIFABPL* gene for *QTgw-3B-12014*. **B**,**C** and **D**, *TaDdRp,TaRLK* and *TaTD* genes for *QGns/Sn/Tgw-4B-1220*, respectively. **E**, *TaTFC3* gene for *QTgw-6A-7089*. **F**, *TaKMT* gene for *QSn/Tgw-7A-9048*. **G**, *TaSPL17* gene for *QGns-7D-6025*.

### *TaDdRp, TaRLK* and *TaTD* genes for *QGns/Sn/Tgw-4B-1220* regulated GNS, GNS and TGW, respectively

The meta-QTL interval of *QGns/Sn/Tgw-4B-1220* for GNS, SN and TGW was at the 1,220.0–1,234.9 cM on chromosome 4B in the UG-Map. This locus included 15 CUGs, including ten CUGs annotated in RefSeq v1.1, two CUGs annotated in the TL-RILs, and three ncRNAs (Fig. 3B; Table 2, S3). Of the eight HC CUGs annotated in RefSeq v1.1. We selected the first CUG, *TraesCS4B02G046500*, and two CUGs in the middle positions of the region, *TraesCS4B02G047100* and *TraesCS4B02G047600*, to confirm their functions.

*TraesCS4B02G046500* is annotated as a DNA-directed RNA polymerase subunit (*TaDdRp-B1* gene) with one InDel in an intron (Table S4, S5). *TaDdRp-B1* was edited and three homozygous mutant genotypes *bb-1* (−11 bp), *bb-2* (+1 bp), *bb-3* (−6 bp) were identified in the T_4_ generation (Fig 4B, S2B). The GNS values of *bb-1, bb-2, bb-3* and WT were 49.14, 44.10, 43.50 and 44.08, respectively. The GNS of *bb-1, bb-2* and *bb-3* were significantly lower than that of WT by 5.04 (10.27%), 5.64 (11.48%) and 5.06 (10.30%), respectively. For SN and TGW, the differences were not significant between the WT and mutants (Fig. 4B; Table S7). The results indicated that the *TaDdRp-B1* gene regulated the GNS.

*TraesCS4B02G047100* is annotated as a receptor-like kinase (*TaRLK-B1* gene) with one non-synonymous substitution (Ser495Leu) in the exon and one SNP in the promoter. The expression levels were significantly different between the TN18 and LM6 genotypes of the TL-RILs (Table S4, S5). *TaRLK-B1* gene was edited, and the three homozygous mutant genotypes *bb-1* (+1 bp), *bb-2* (−2 bp) and *bb-3* (−1 bp) were identified in the T_4_ generation (Fig. 4C, S2C). The GNS values of *bb-1, bb-2, bb-3* and WT were 42.31, 41.32, 40.29 and 47.34, respectively. The GNS of *bb-1, bb-2* and *bb-3* were significantly lower than that of WT by 5.03 (10.63%), 6.02 (14.23%) and 7.06 (17.08%), respectively. For SN and TGW, the differences were not significant between the WT and mutants (Fig. 4C; Table S8). The results indicated that the *TaRLK-B1* gene regulated the GNS.

*TraesCS4B02G047600* is annotated as a threonine dehydratase (*TaTD-B1* gene) with one stop-gain (2,204 bp from the start site of the 5’UTR) and one non-synonymous substitution (Glu573Lys) in the exon and no SNP/InDel in the promoter. The expression levels were significantly different between the TN18 and LM6 genotypes of the TL-RILs (Table S4, S5). The three homologous genes of *TaTD* on subgenomes A (*TraesCS4A02G266700*), B and D (*TraesCS4D02G047500*) were edited simultaneously. Three homozygous mutant genotypes, *aa* (subgenome A, -27 bp), *bb* (−17 bp) and *dd* (subgenome D, -1 bp), were identified in the T_4_ generation (Fig. 4D, S2D). The TGW values of *aa, bb, dd* and WT were 45.39, 45.65, 45.64 and 40.96 g, respectively. The TGW of mutants *aa, bb* and *dd* were significantly higher than that of WT by 4.44 (10.83%), 4.70 (11.47%) and 4.68 g (11.43%), respectively. For SN and GNS, the differences were not significant between the WT and mutants (Fig. 4D; Table S9). The results indicated that the *TaTD-A1, TaTD-B1* and *TaTD-D1* genes regulated TGW.

### *TaTFC3* gene for *QTgw-6A-7089* regulated TGW

The meta-QTL interval of *QTgw-6A-7089* for TGW was at the 7,086.8-7,092.3 cM on chromosome 6A in the UPG-Map. This locus contained only one CUG, *TraesCS6A02G221300* (Fig. 3C; Table 2, S3). *TraesCS6A02G221300* is annotated as a B-block binding subunit of TFIIIC (*TaTFC3-A1*) with two SNPs in introns. The expression levels were significantly different between the TN18 and LM6 genotypes of the TL-RILs (Table S4, S5). We performed gene editing of *TaTFC3-A1* and two homozygous mutant genotypes, *aa-1* (−2 bp) and *aa-2* (+1 bp), were identified in T_4_ generation (Fig. 4E, S2E). The TGW values of *aa-1, aa-2* and WT were 50.97, 52.66 and 45.37 g, respectively. The TGW of mutants *aa-1*and *aa-2* were significantly higher than that of WT by 5.60 (12.34%) and 7.29 g (16.01%), respectively (Fig. 4E; Table S10). The results indicated that *TaTFC3-A1* gene regulated TGW.

### *TaKMT* gene for *QSn/Tgw-7A-9048* regulated SN

The meta-QTL interval of *QSn/Tgw-7A-9048* for SN and TGW was at the 9,037.5–9,069.5 cM on chromosome 7A in the UG-Map. This region involved only one CUG, *TraesCS7A02G364700* (Fig. 3D; Table 2, S3). *TraesCS7A02G364700* is annotated as a histone-lysine N-methyltransferase (*TaKMT-A1* gene) with four SNPs and one InDel in the 5’UTR and no SNP/InDel in the promoter. The expression levels were significantly different between the TN18 and LM6 genotypes of the TL-RILs (Table S4, S5). We performed gene editing of *TaKMT-A1*. Two homozygous mutant genotypes, *aa-1* (−1 bp) and *aa-2* (−2 bp), were identified in the T_4_ generation (Fig. 4F, S2F). The SN values of *aa-1, aa-2* and WT were 3.43, 3.48 and 4.48, respectively. The SN of mutants *aa-1* and *aa-2* were significantly lower than that of WT by 1.06 (23.54%) and 1.01 (22.43%), respectively. For TGW, the differences were not significant between the WT and mutants (Fig. 4F; Table S11). The results indicated that the *TaKMT-A1* gene regulated SN.

### *TaSPL17* gene for *QGns-7D-6025* regulated GNS

The meta-QTL interval of *QGns-7D-6025* for GNS was at the 6,013.0–6,026.0 cM on chromosome 7D in the UG-Map. This locus involved four CUGs with two HC CUGs, *TraesCS7D02G245100* and *TraesCS7D02G245200* (Fig. 3E; Table 2, S3). One SNP at 213,787,168 bp was found in the sequences of the two HC CUGs. For *TraesCS7D02G245100*, the SNP is in the 3’UTR. For *TraesCS7D02G245200*, the SNP causes a non-synonymous substitution (Asn356Tyr) in the exon (Table S4). Therefore, we selected *TraesCS7D02G245200* to confirm its function. *TraesCS7D02G245200* is annotated as a squamosa promoter-binding-like protein 17 (*TaSPL17-D1* gene). The three homologous genes on subgenomes A (*TraesCS7A02G2465000*), B (*TraesCS7B02G144900*) and D of *TaSPL17* were edited simultaneously. Three homozygous mutant genotypes, *aa* (−1 bp), *dd-1* (+1 bp) and *dd-2* (−2 bp), were identified in the T_4_ generation (Fig 4G, S2G). The GNS values of *aa* and WT were 41.15 and 47.23, respectively. The GNS values of *dd-1, dd-2* and WT were 36.40, 35.07 and 43.10, respectively. The GNS of mutants *aa, dd-1* and *dd-2* were significantly lower than that of WT by 6.07 (12.86%), 6.70 (15.55%) and 8.03 (18.63%), respectively (Fig. 4G; Table S12). The results indicated that the *TaSPL17-A1* and *TaSPL17-D1* genes regulated GNS.

## Discussion

### Previously cloned orthologous genes of CUGs

The CUGs for which orthologous genes have been cloned previously are highly reliable and their functions should be confirmed preliminarily (Vercruysse et al. 2019). For the 75 HC CUGs annotated in RefSeq v1.1, six orthologous genes for five CUGs were previously cloned with the same or similar agronomic functions (Table S13). For GNS related traits, four orthologous genes for three CUGs were obtained. *OsUCL8* of rice was the orthologous gene of *TraesCS4B02G044600* for *QGns/Sn-4B-1132. OsUCL8* is an *uclacyanin* (*UCL*) gene of the phytocyanin family. The knock down or knock out of *OsUCL8* increases grain yield, while the overexpression of *OsUCL8* results in an opposite phenotype (Zhang *et al*., 2017). *ADL1C* and *DRP1A* of *Arabidopsis* were the orthologous genes of *TraesCS4B02G047700* for *QGns/Sn/Tgw-4B-1220. ADL1C*, a member of the *Arabidopsis dynamin-like protein* (*ADL*) family, is essential for the formation and maintenance of the pollen cell surface and viability during desiccation (Kang et al., 2003). Moreover, a null mutant of the *dynamin-related protein 1A* (*DRR1A*), *rsw9*, reduced the final heights and resembled flowers of *adl1a* in that stigma papillae failed to elongate and seed set was low (Kang et al., 2003). But seeds that did form in *rsw9* were not shrivelled as found for *adl1a* seed (Kang et al., 2001). *OsSPL14* (*squamosa promoter binding protein-like 14, IPA1*) of rice was the orthologous gene of *TraesCS7D02G245200* for *QGns-7D-6025*. Higher expression of *OsSPL14* in the reproductive stage promotes panicle branching and higher grain yield in rice. *OsSPL14* controls shoot branching in the vegetative stage (Miura *et al*., 2010).

For TGW related traits, one orthologous genes for one CUG was found. *GSD1* (*Grain setting defect1*) of rice was the orthologous gene of *TraesCS2D02G464900* for *QTgw-2D-6108. GSD1* encodes a putative remorin protein. A mutant *gsd1-D* was reduceds the grain setting rate, 1,000-grain weight and grain thickness (Gui et al., 2014).

For SN related traits, one orthologous genes was found. *COL7* (*CONSTANS-LIKE 7*) of *Arabidopsis* was the orthologous gene of *TraesCS4B02G045700* for *QSn-4B-1170*. Overexpression of *COL7* enhances branching number only when plants are grown in regular white light but not in shade via the suppression of auxin level in a phyB-dependent manner (Wang et al., 2013; Zhang et al., 2014).

### Efficiency of QTL-Seq-RIL

We previously cloned a reduced-height (*Rht*) gene *TaOSCA1*.*4* from a QTL *QPh-1B* in the “Chuan35050 × Shannong483” RIL population. *QPh-1B* included an EST-SSR marker *swes1079*, which represent a partial sequence of a gene. We sequenced the *TaOSCA1*.*4*, which coantained *swes1079*, and validated its function via RNAi technology (Lv et al., 2024). This inspires us that cloning genes from QTLs of RIL populations is feasible. So we proposed the QTL-Seq-RIL for cloning genes. Uing QTL-Seq-RIL, we identified 138 CUGs for YCTs from 77 QTLs of a single RIL population, and the functions of seven CUGs, *TaIFABPL, TaDdRp, TaRLK, TaTD, TaTFC3, TaKMT* and *TaSPL17*, were confirmed via the CRISPR/Cas9 system. Moreover, the orthologous genes of five CUGs (include *TaSPL17*) with the same or similar agronomic functions were previously cloned and their functions should be preliminarily validated (Vercruysse et al., 2019). It is to say, 11 of the 75 HC CUGs (14.7%) from a single RIL population were preliminarily confirmed. The other CUGs need to be further confirmation. In addition, many genes could be cloned for other traits using RIL populations. For example, we have cloned a quality gene *XIP (Xylanase Inhibitor Protein)* (Sun et al., 2022) and a *Rht* gene *TaDHL* (*ATP-dependent DNA helicase*) (Guo et al., 2022) from QTLs of the TL-RILs. Seven other *Rht* genes were also confirmed via CRISPR/Cas9 technology from QTLs of the TL-RILs and “Shannong0431 × Lumai21” RILs (unpublished data). In wheat, hundreds of RIL populations have been constructed, and their tremendous corresponding phenotypic data have been investigated (Cao et al., 2020; Saini et al., 2020). QTL-Seq-RIL provides an efficient method for rapid gene cloning using existing RILs.

### Key point of QTL-Seq-RIL

Using the RIL population to construct the UG-Map is the key point of the QTL-Seq-RIL strategy. Firstly, the RIL population, originating from F_2_ individuals through repeated self-crossing, accumulates crossover events over generations. This results in a significant increase in crossover frequency and, consequently, extensive recombination and greater distances between adjacent genes. (Fig. S1; Table S1). This high level of recombination in advanced RIL generations makes them ideally suited for accurate QTL localization, requiring only hundreds of lines compared to the thousands typically needed for map-based cloning. Secondly, the UG-Map was constructed based on physical positions of all unigenes in the reference genome, without omitting any unigenes. This method aligns with the fundamental concept of a genetic linkage map, providing a more accurate gene order than traditional genetic mapping using DNA markers. We also constructed the genetic map for sub-unigenes of the TL-RILs using the current mapping motheds of markers. It showed a high ratio of un-linked sub-unigenes (Table S14) and serious false linkages between sub-unigenes (Table S15), which enormously affecting the comprehensiveness and accuracy of QTL mapping. Thirdly, using the RNA-Seq technology, we can directly detect the polymorphic unigenes in whole genome from a RIL population with much lower cost than that of DNA sequencing (wheat transcriptome, ∼500 Mb; genome, ∼17 Gb). We also sequenced the DNA for a subset of RILs and thier parents to determine the polymorphism of the promoter (Table S5). This dual approach offers a more comprehensive understanding of candidate genes. In addition, we used multi-environment trials and three QTL mapping software programmes to reduce false positive QTLs. As a result of all these factors, we identified an average of 1.8 candidate genes per QTL using UG-Map (Table 2). This number was ultra-smaller, which enabled us to confirm the CUGs directly with less effort and higher efficiency.

### Potential use of the cloned genes in wheat yield improvement

Of the seven genes we cloned in this study, no orthologous genes for *TaIFABPL, TaDdRp, TaRLK, TaTD, TaTFC3* and *TaKMT* were previously cloned with the same or similar agronomic functions. These six genes were first found to regulate YCTs in crops. *TaSPL17* is the orthologous gene of *OsSPL14*, which controls panicle branching and grain yield (Miura *et al*., 2010).

In the TL-RILs, the haplotypes of the coned genes, with a few polymorphic sites, showed significant differences on phenotype using *t*-test (Table S16). *TaRLK-B1, TaTD-B1* and *TaSPL17-D1* had one, two and one missense mutation(s) in the exons, respectively (Table S4). For *TaRLK-B1*, TN18 type significantly increased the GNS of 13.82, 8.92. 7.98, 6.49 and 7.93% under F(AV), M(CKAV), M(LNAV), M(LPAV) and M(LKAV) environments (five AV environments), respectively, comparing with LM6 type. For *TaTD-B1*, TN18 type significantly increased the TGW of 8.48, 3.63. 3.35, 3.07 and 3.46% under the five AV environments, respectively. For *TaSPL17-D1*, TN18 type increased the GNS of 5.82, 4.88. 3.63, 3.57 and 3.53% under the five AV environments, respectively. *TaIFABPL-A1* and *TaKMT-A1* had one and four mutation in the UTR, respectively (Table S4). LM6 type of *TaIFABPL-A1* significantly increased TGW of 6.87, 2.91. 3.51and 3.30% under the five AV environments, respectively. LM6 type of *TaKMT-A1* significantly increased SN of 16.26, 12.53. 10.59, 11.28 and 11.89% under the five AV environments, respectively. *TaDdRp-B1* and *TaTFC3-A1* had one and two mutation(s) in the intron, respectively (Table S4). For *TaDdRp-B1*, TN18 type significantly increased the GNS of 15.19, 8.50. 10.03, 6.39 and 8.33% under the five AV environments, respectively. For *TaTFC3-A1*, TN18 type significantly increased the TGW of 7.81, 3.35. 5.65, 4.81 and 4.47% under the five AV environments, respectively. In yeast, both synonymous and nonsynonymous mutations affected the fitness effects, which negated the presumption that synonymous mutations are neutral or nearly neutral (Shen et al., 2022). It may be a reason that the mutation(s) in the intron can cause the phenotypic variations. These results indicated that the TN18 haplotypes of *TaDdRp-B1, TaRLK-B1, TaTD-B1, TaTFC3-A1* and *TaSPL17-D1*, and LM6 haplotype of *TaIFABPL-A1* and *TaKMT-A1* are favourable in improving grain yield, especially under field environments. Due to TN18 and LM6 are varieties developed in recent years, the haplotypes of the four genes can found in current varieties, which are the important parents. Furthermore, TN18 and its derived varieties are core parents in the Huang Huai Wheat Region, China. These genes should play an important role in present wheat yield improvement.

Interestingly, the knockout mutants of *TaIFABPL, TaTFC3-A1, TaTD-A1, TaTD-B1* and *TaTD-4D1* increased the TGW by 9.10–16.01%. Of these, *TaTD-A1* and *TaTD-4D1* were novel mutants which were not found in the TL-RILs and present varieties.

## Materials and Methods

### Plant materials

The plant materials were a set of RILs derived from a cross of ‘Tainong 18 × Linmai 6’ (TN18 × LM6, TL-RILs, F_11_ in the 2015–2016 growing season) by single-seed descent (SSD). A total of 184 lines randomly selected from the original 305 lines were used to conduct field trials and RNA-Seq analysis. TN18 is a cultivar developed by our group that was authorized for commercial release in 2008 by the Crop Variety Approval Committee (CVAC) of Shandong Province, China. It is a popular cultivar and has become a core parent from which over 60 authorized cultivars had been developed as of 2023. LM6 is an elite breeding line developed by the Linyi Academy of Agricultural Sciences, Shandong, China. ‘TN18 × LM6’ is an excellent cross for wheat breeding programs. We developed two cultivars, Shannong 29 and Shannong 41, from this cross and authorized by the National CVAC of China in 2016 and 2021, respectively. Another cultivar, Shannong 30, was developed from reciprocal crosses of the TL-RILs and authorized in 2017 by the National CVAC of China. Shannong 29 was planted 500 thousand hectares in 2021, which was the fifth rank in area of the wheat cultivars in China.

### RNA and DNA extraction and sequencing

The samples used for RNA extraction consisted of five plants of each line of the TL-RILs and the parents from the field trial of the 2015–2016 growing season at seven stages in the process of wheat growth, including the seedling establishment, jointing, flag leaf, flowering, grain-filling I (10 days after flowering stage), grain-filling II (20 days after flowering stage) and grain-filling III (30 days after flowering stage) stages. In the first three stages, one tiller of each plant was sampled; while in the last four stages, one spike of each plant was sampled. Total RNA was extracted from frozen tissue using the RNAprep Pure Plant Kit (TIAGEN, Beijing, China) and the RNAprep Pure Plant Kit (Polysaccharides-& Polyphenolics-rich) (TIAGEN, Beijing, China). For each of the parents, RNA extraction was conducted in each of the seven stages. For the RIL population, RNA extraction was conducted using a mixture of samples from the first three stages and a mixture of samples from the last four stages. Frozen tissue was ground with a mortar, and approximately 100 mg of powdered tissue was sampled and used for RNA extraction following the kit manufacturer’s instructions. RNA concentrations were measured using a Qubit RNA Assay Kit on a Qubit 2.0 Fluorometer (Life Technologies, CA, USA). RNA purity was checked using a Nanophotometer spectrophotometer (IMPLEN, CA, USA). RNA integrity was assessed using the RNA Nano 6000 Assay Kit for the Agilent Bioanalyzer 2100 system (Agilent Technologies, CA, USA). RNA samples were stored at -80 °C until being sent for sequencing.

cDNA libraries with different insert sizes were constructed using the NEB Next Ultra TM Directional RNA Library Prep Kit for Illumina (NEB, USA) following the manufacturer’s recommendations and were assessed on an Agilent Bioanalyzer 2100 system. A total of 198 cDNA libraries (184 from the first three stages of the RILs and all seven stages of the parents TN18 and LM6) and 191 cDNA libraries (184 from the last four stages of the RILs and all seven stages of the TN18 parent) were sequenced using an Illumina HiSeq X Ten sequencer by Annoroad Gene Technology Co. (Beijing, China) and Novogene Bioinformatics Institute (Beijing, China), respectively. Thus, 150 bp paired-end reads were generated, and approximately 12 Gb (∼30×) of clean paired-end reads from each library were obtained from the two companies.

To identify the mutants in promoters of unigenes, twenty samples, including samples of TN18, LM6 and 18 RILs, were used for DNA sequencing. Seeds were sown on absorbent paper and grown to the two leaf stage, and DNA was then extracted using the Hi-DNAsecure Plant Kit (TIAGEN, Beijing, China). Qualified DNA fragmentation was carried out with an ultrasonic processor, and a 350 bp library was prepared after terminal end repair, sequence adapter addition and purification. The library insert size was verified with an Agilent 2100 system. Finally, Q-PCR was implemented to ensure the effective quantified library concentrations. Twenty samples were sequenced on an Illumina HiSeq X Ten machine using 150-base pair paired-end reads by Annoroad Gene Technology Co. (Beijing, China) with approximately 450 Gb (∼30×) and 150 Gb (∼10×) of clean paired-end reads for each parent and each line of RILs, respectively.

### Assembly of unigenes, SNP/InDel calling and annotation of SNPs/InDels to unigenes

For both RNA-Seq and DNA-Seq, the raw reads were trimmed using Trimmomatic (Bolger et al., 2014), and high-quality clean reads were mapped to the RefSeq v1.1 reference genome (IWGSC, 2018) using HISAT2 (Kim et al., 2015). Only reads that were uniquely mapped to the genome were retained for further steps. The sorting of BAM files, duplicate removal and read group addition were performed with Picard Tools 2.9.0 (http://broadinstitute.github.io/picard/). SNPs/InDels were identified with the HaplotypeCaller module of GATK v3.8 in GVCF mode. Then, joint calling was performed using GATK GenotypeGVCFs. SNPs were preliminarily filtered using the GATK VariantFiltration function with the following parameters: QD < 2.0, FS > 60.0, MQ < 40.0, MQ rank sum < -12.5, read post rank sum < -8.0, cluster window size 35, and cluster size 3. InDels were filtered using the following parameters: QD < 2.0, FS > 200.0, read post rank sum < -20.0, cluster window size 35, cluster size 3. SNPs/InDels with more than 80% missing data or a minor allele frequency (MAF) below 2.5% were discarded (Mascher et al., 2013). Heterozygous calls among the RILs were set to missing. Furthermore, the HISAT2-StringTie pipeline was used to assemble the parents and 80 RILs as the supplemental annotation of RefSeq v1.1. The above software was run mainly using a Tianhe-2 supercomputer rented from the National Supercomputer Center in Guangzhou, China, and computer server of the State Key Laboratory of Crop Biology, Shandong Agricultural University.

Using a Python script, the SNPs/InDels were annotated to three types of unigenes: RefSeq genes, TL-RIL genes and ncRNAs. For the TL-RIL genes, we used the hisat2-stringtie pipeline to annotate genes. Augustus was used to predict gene models with the command ‘Augustus – species = wheat –strand = both – singlestrand = false – genemodel = partial – codingseq = on – sample = 6769 –keep_viterbi = true – alternatives-from-sampling = true – minexonintronprob = 0.2 –minmeanexonintronprob = 0.5 – maxtracks = 2’ (Stanke and Waack, 2003). The genes predicted via the above steps were considered to be the TL-RIL genes, and the rests were considered to be ncRNAs.

### Construction of genetic map of unigenes (UP-Map)

The UG-Map was constructed as follows: 1) Division of unigenes into sub-unigenes. When a unigene included two or more SNPs/InDels, the genetic distances between the SNPs/InDels were calculated using IciMapping 4.1 (http://www.isbreeding.net) (Meng et al., 2015). If the genetic interval between two physically adjacent markers was larger than 5 cM, they were divided into different sub-unigenes. 2) Genotyping of sub-unigenes. The sub-unigene genotypes were merged to obtain one genotype using MSTMap (Wu et al., 2008) in the R language with the following parameters: method = “maxmarginal”, error.prob = 0.01, map.function = “Kosambi”. 3) Filtering genotypes of sub-unigenes. All of the sub-unigenes that showed an MAF greater than 5% and missing data less than 80% of the TL-RILs were reserved (Mascher et al., 2013). Furthermore, if a unigene contained more than five sub-unigenes, only five sub-unigenes with the least missing data were retained. 4) Construction of UG-Map. The sub-unigenes were sorted by physical position according to RefSeq v1.1(IWGSC, 2018) and employed to construct an UG-Map using MapMaker software (Lander et al., 1987). When a genetic distance was more than 20 cM, we noted as 20 cM.

### Design of trials and measurement of traits

The field trials (noted as F) were conducted in six growing seasons (environments), in 2010–2011 (F(E11)), 2011– 2012 (F(E12)), 2012–2013 (F(E13)), 2013–2014 (F(E14)), 2014–2015 (F(E15)) and 2015–2016 (F(E16)), at the Experimental Station of Shandong Agricultural University (Tai’an, China). Average values (AV) were computed and are noted as F(AV). Seeds were sown on October 5–10, and plants were harvested on June 10–15 of the next year. Each plot consisted of three rows, which were 1.5 m long and spaced 25 cm apart, with two repetitions. Fifty seeds were planted in each row.

Mineral nutrient trials in the maturation stage (noted as M) were carried out in eight 110 m^2^ (10 × 11 m) nutrient pools. We constructed the nutrient pools in 2008 at the Experimental Station of Shandong Agricultural University (Tai’an, China) to perform a whole-growth-stage trial of mineral nutrient elements. The nutrient pools were segregated using a cement brick wall with a 1.5 m depth. The soil structure of a natural field with loamy soil was maintained. The mineral nutrient elements were consumed by annually planting wheat and maize until the nutrient contents were in accordance with the requirements of the trials (Kong et al., 2013). The nutrient pool trials were conducted in four growing seasons (environments), 2012–2013, 2013–2014, 2014–2015 and 2015–2016, with two replications in each season. Four treatments were set up: normal mineral nutrition (CK, grain yield: 9000 kg/ha) and low N, P and K nutrition (LN, LP and LK; grain yield: 6000 kg/ha for each corresponding element). The environments/treatments were noted as M(CK13), M(CK14), M(CK15), M(CK16), M(CKAV), etc. Seeds were sown on October 10–15, and plants were harvested on June 10–15 the next year. In 2012–2013 and 2013–2014, twenty seeds of each line were sown per 1 m row, with spacing of 5 cm between plants and 25 cm between rows. In 2014– 2015 and 2015–2016, forty seeds of each line were sown per row, with spacing of 2.5 cm between plants and 25 cm between rows.

The GNS were measured in 10 randomly sampled plants in each replication, SN was measured the spike number in a 50 cm length within one row and then counting the spike number per m^2^, and TGW was measured by weighing 200 grains three times after harvest.

### QTL analysis, candidate unigenes (CUGs) identification and orthologous gene acquisition

The AVs were taken as environments. Trait values for each line of the TL-RILs in the 27 environments/treatments were used for QTL analysis by Windows QTL Cartographer 2.5 (http://statgen.ncsu.edu/qtlcart/WQTLCart.htm), IciMapping 4.1 (http://www.isbreeding.net/) and MapQTL 6.0 software (Van Ooijen, 2009). For Windows QTL Cartographer 2.5, composite-interval mapping (CIM) was selected to search for QTLs. The parameter setup ‘model 6 standard analysis’ was used, with a walk speed of 0.5 cM, ‘forward and backward’ regression for the selection of the markers to control for the genetic background, up to five control markers, and a blocked window size of 10 cM to exclude closely linked control markers at the testing site. For IciMapping 4.1, inclusive composite interval mapping (ICIM) (Li et al., 2007) was carried out with a step size of 0.5 cM. The parameter for handling missing phenotypic data was ‘Deletion’. The largest *P*-value for entering variables in the stepwise regression of phenotype on marker variables was 0.001. For MapQTL 6.0, the multiple-QTL model (MQM) package with a mapping step size of 0.5 cM was used to map QTLs. The ‘Regression’ algorithm was selected. The maximum number of neighboring markers was 5, the functional tolerance value was 1.0E-08, and the number of permutations was 1000. The LOD threshold for declaring a significant QTL in the three software programs was an LOD>2.5, with an LOD≥3.0 for at least one environment (He et al., 2020; Li et al., 2021; Xu et al., 2021). The QTL boundaries were determined using a meta-QTL software of BioMercatorV3.0 (Sosnowski et al., 2012).

For the field trials, we defined a stable QTL for a single software as a QTL that was detected in AV and more than two environments. For the mineral nutrient trials, we defined a stable QTL as a QTL detected in in AV and more than one environment/treatment for CK, LN, LP or LK. A stable QTL was declared to exist when it was detected by at least two of the three software, Windows QTL Cartographer 2.5, IciMapping 4.1 and MapQTL 6.0. The unigenes covered by the interval of QTLs were regarded as the CUGs of the corresponding QTLs.

We identified the rice and *Arabidopsis* orthologous genes of the wheat genes using ortholog information downloaded via BioMart (http://plants.ensembl.org/biomart/martview) (Kersey et al., 2018) on Ensembl Plants Genes 51. Functional annotations were obtained from funricegenes (Yao et al., 2018), RAP-DB (Sakai et al., 2013), Araport (Cheng et al., 2017), and literature searches. The R package Biostrings (Pagès et al., 2019) were also used, in which the ‘pairwise Alignment’ function to perform a ‘global’ alignment of protein sequences and then the ‘pid’ function to calculate the percent sequence identity. If genes between wheat and other species had the alignment score greater than 0 and the protein sequence similarity over 50%, they were assumed to have similar functions.

### Gene editing by the CRISPR/Cas9 system

To identify the function of CUGs in wheat, we knocked out corresponding genes using the CRISPR/Cas9 system. The genes of *TaIFABPL-B1, TaDdRp-B1, TaRLK-B1, TaTFC3-A1* and *TaKMT-A1* were knocked out using one sgRNA. For *TaTD* and *TaSPL17* genes, three homogenous genes on subgenomes A, B and D were knocked out using the same two sgRNAs. The sequences of the genes were obtained from the Fielder genome and used to design sgRNA target sequences in CRISPR-direct (http://crispr.dbcls.jp/) and CRISPOR (http://crispor.tefor.net/). The gene editing was conducted by the Crop Research Institute, Shandong Academy of Agricultural Sciences, China (Zhang et al., 2018). Sequence mutations were detected in the T1-T4 generations via Hi-TOM analysis (Liu et al., 2019). For T_4_ generation, twenty seeds of mutant genotypes and WT were sown per 1 m row, with spacing of 5 cm between plants and 25 cm between rows. One row of WT was planted between every two rows of mutants.

## Acknowledgements

This work was supported by the National Key Research and Development Programme of China (2021YFD1200600), the Agricultural Variety Programme of Shandong Province, China (2021LZGC013 and 2022LZGCQY002) and the Modern Agricultural Industry Technology System of Shandong Province, China.

## Author contributions

S.L., Y.G. and Y.Y. conceived the study and designed the experiment. M.Z., X.H., H.W., J.S., B.G., M.G. H.X,, G.Z., H.L., X.C., N.L., Y.X., Q.W., C.W., G.Z., Y.Y., J.M. Y.P., G.L., C.Q., J.S., X. C., X. D., F.K., Y.Z. and Y.A. performed the experiments and data analysis. M.Z., S.L. and B.G.wrote the manuscript. All authors read and approved the final manuscript.

## Data availability

All data supporting the findings of this study are available within the paper and within the supplemental data published online or can reasonably request from the corresponding author.

## References

Arora S, Steuernagel B, Gaurav K, Chandramohan S, Long Y, Matny O, Johnson R, Enk J, Periyannan S, Singh N et al. 2019. Resistance gene cloning from a wild crop relative by sequence capture and association genetics. Nature Biotechnology 37: 139–143.:

Athiyannan N, Abrouk M, Boshoff WHP, Cauet S, Rodde N, Kudrna D, Mohammed N, Bettgenhaeuser J, Botha KS, Derman SS et al. 2022. Long-read genome sequencing of bread wheat facilitates disease resistance gene cloning. Nature Genetics 54: 227–231.

Bolger AM, Lohse M, Usadel B. 2014. Trimmomatic: a flexible trimmer for Illumina sequence data. Bioinformatics 30: 2114–2120.

Cao S, Xu D, Hanif M, Xia X, He Z. 2020. Genetic architecture underpinning yield component traits in wheat. Theoretical and Applied Genetics 133: 1811–1823.

Chen Y, Yan Y, Wu TT, Zhang GL, Yin H, Chen W, Wang S, Chang F, Gou JY. 2020. Cloning of wheat keto-acyl thiolase 2B reveals a role of jasmonic acid in grain weight determination. Nature Communications 11: 6266.

Chen Z, Ke W, He F, Chai L, Cheng X, Xu H, Wang X, Du D, Zhao Y, Chen X et al. 2022. A single nucleotide deletion in the third exon of FT-D1 increases the spikelet number and delays heading date in wheat (Triticum aestivum L.). Plant Biotechnology Journal 20: 920–933.

Cheng CY, Krishnakumar V, Chan AP, Thibaud-Nissen F, Schobel S, Town CD. 2017. Araport11: a complete reannotation of the Arabidopsis thaliana reference genome. Plant Journal 89: 789–804.

Cheng X, Xin M, Xu R, Chen Z, Cai W, Chai L, Xu H, Jia L, Feng Z, Wang Z et al. 2020. A single amino acid substitution in STKc_GSK3 kinase conferring semispherical grains and its implications for the origin of Triticum sphaerococcum Perc. Plant Cell 32: 923–934.

Choulet F, Alberti A, Theil S, Glover N, Barbe V, Daron J, Pingault L, Sourdille P, Couloux A, Paux E et al. 2014. Structural and functional partitioning of bread wheat chromosome 3B. Science 345: 1249721.

Dong C, Zhang L, Zhang Q, Yang Y, Li D, Xie Z, Cui G, Chen Y, Wu L, Li Z et al. 2023. Tiller Number1 encodes an ankyrin repeat protein that controls tillering in bread wheat. Nature Communications 14: 836.

Du D, Zhang D, Yuan J, Feng M, Li Z, Wang Z, Zhang Z, Li X, Ke W, Li R et al. 2021. FRIZZY PANICLE defines a regulatory hub for simultaneously controlling spikelet formation and awn elongation in bread wheat. New Phytologist 231: 814–833.

Feuillet C, Travella S, Stein N, Albar L, Nublat A, Keller B. 2003. Map-based isolation of the leaf rust disease resistance gene Lr10 from the hexaploid wheat (Triticum aestivum L.) genome. Proceedings of the National Academy of Sciences of the United States of America 100: 15253–15258.

Fu D, Uauy C, Distelfeld A, Blechl A, Epstein L, Chen X, Sela H, Fahima T, Dubcovsky J. 2009. A kinase-START gene confers temperature dependent resistance to wheat stripe rust. Science 323: 1357–1360.

Gui J, Liu C, Shen J, Li L. 2014. Grain setting defect1 encoding a remorin protein affects the grain setting in rice through regulating plasmodesmatal conductance. Plant Physiology 166: 1463–1478.

Guo B, Jin X, Chen J, Xu H, Zhang M, Lu X, Wu R, Zhao Y, Guo Y, An Y, Li S. 2022. ATP-dependent DNA helicase (TaDHL), a novel reduced-height (Rht) gene in wheat. Genes 13: 979.

Godfray HCJ, Beddington JR, Crute IR, Haddad L, Lawrence D, Muir JF, Pretty J, Robinson S, Thomas SM, Toulmin C. 2010. Food security: the challenge of feeding 9 billion people. Science 327: 812–818.

He X, Kabir MR, Roy KK, Anwar MB, Xu K, Marza F, Odilbekov F, Chawade A, Duveiller E, Huttner E et al. 2020. QTL mapping for field resistance to wheat blast in the Caninde#1/Alondra population. Theoretical and Applied Genetics 133: 2673–2683.

IWGSC. 2014. A chromosome-based draft sequence of the hexaploid bread wheat (Triticum aestivum) genome. Science 345: 1251788.

IWGSC. 2018. Shifting the limits in wheat research and breeding using a fully annotated reference genome. Science 361: eaar7191.

Kang BH, Busse JS, Dickey C, Rancour DM, Bednarek SY. 2001. The Arabidopsis cell plate-associated dynamin-like protein ADL1Ap is required for multiple stages of plant growth and development. Plant Physiology 126: 47–68.

Kang BH, Rancour DM, Bednarek SY. 2003. The dynamin-like protein ADL1C is essential for plasma membrane maintenance during pollen maturation. Plant Journal 35: 1–15.

Kersey PJ, Allen JE, Allot A, Barba M, Boddu S, Bolt BJ, Carvalho-Silva D, Christensen M, Davis P, Grabmueller C et al. 2018. Ensembl Genomes 2018: an integrated omics infrastructure for non-vertebrate species. Nucleic Acids Research 46(D1): D802–D808.

Kim D, Langmead B, Salzberg SL. 2015. HISAT: a fast spliced aligner with low memory requirements. Nature Methods 12: 357–360.

Kong FM, Guo Y, Liang X, Wu CH, Wang YY, Zhao Y, Li SS. 2013. Potassium (K) effects and QTL mapping for K efficiency traits at seedling and adult stages in wheat. Plant Soil 373: 877–892.

Lander ES, Green P, Abrahamson J, Barlow A, Daly MJ, Lincoln SE, Newberg LA. 1987. MAPMAKER: an interactive computer package for constructing primary genetic linkage maps of experimental and natural populations. Genomics 1: 174–181.

Li G, Zhou J, Jia H, Gao Z, Fan M, Luo Y, Zhao P, Xue S, Li N, Yuan Y et al. 2019. Mutation of a histidine-rich calcium-binding-protein gene in wheat confers resistance to Fusarium head blight. Nature Genetics 51: 1106–1112.

Li H, Ye G, Wang J. 2007. A modified algorithm for the improvement of composite interval mapping. Genetics 175: 361–374.

Liu Q, Wang C, Jiao X, Zhang H, Song L, Li Y, Gao C, Wang K. 2019. Hi-TOM: a platform for high-throughput tracking of mutations induced by CRISPR/Cas systems. Science China-Life Sciences 62: 1–7.

Lu P, Guo L, Wang Z, Li B, Li J, Li Y, Qiu D, Shi W, Yang L, Wang N et al. 2020. A rare gain of function mutation in a wheat tandem kinase confers resistance to powdery mildew. Nature Communications 11: 680.

Li T, Deng G, Su Y, Yang Z, Tang Y, Wang J, Qiu X, Pu X, Li J, Liu Z et al. 2021. Identification and validation of two major QTLs for spike compactness and length in bread wheat (Triticum aestivum L.) showing pleiotropic effects on yield-related traits. Theoretical and Applied Genetics 134: 3625–3641.

Lv G, Jin X, Wang H, Wang Y, Wu Q, Wu Jiang F, Ma Y, An Y, Zhang M. et al. 2024. Cloning a novel reduced-height (Rht) gene TaOSCA1.4 from a QTL in wheat. Frontiers in Plant Science (under review)

Mascher M, Muehlbauer GJ, Rokhsar DS, Chapman J, Schmutz J, Barry K, Muñoz-Amatriaín M, Close TJ, Wise RP, Schulman AH et al. 2013. Anchoring and ordering NGS contig assemblies by population sequencing (POPSEQ). Plant Journal 76: 718–727.

Meng L, Li H, Zhang L, Wang J. 2015. QTL IciMapping: integrated software for genetic linkage map construction and quantitative trait locus mapping in biparental populations. Crop Journal 3: 269–283.

Miura K, Ikeda M, Matsubara A, Song XJ, Ito M, Asano K, Matsuoka M, Kitano H, Ashikari M. 2010. OsSPL14 promotes panicle branching and higher grain productivity in rice. Nature Genetics 42: 545–549.

Ni F, Qi J, Hao Q, Lyu B, Luo MC, Wang Y, Chen F, Wang S, Zhang C, Epstein L et al. 2017. Wheat Ms2 encodes for an orphan protein that confers male sterility in grass species. Nature Communications 8: 15121.

Pagès H, Aboyoun P, Gentleman R, DebRoy S. 2019. Biostrings: efficient manipulation of biological strings. R package version 2.52.0.

Periyannan S, Moore J, Ayliffe M, Bansal U, Wang X, Huang L, Deal K, Luo M, Kong X, Bariana H et al. 2013. The gene Sr33 an ortholog of barley Mla genes encodes resistance to wheat stem rust race Ug99. Science 341: 786–788.

Saini DK, Devi P, Kaushik P. 2020. Advances in genomic interventions for wheat biofortification, a review. Agronomy 10: 62.

Sakai H, Lee SS, Tanaka T, Numa H, Kim J, Kawahara Y, Wakimoto H, Yang C, Iwamoto M, Abe T et al. 2013. Rice annotation project database (RAP-DB): an integrative and interactive database for rice genomics. Plant and Cell Physiology 54: e6.

Sakuma S, Golan G, Guo Z, Ogawa T, Tagiri A, Sugimoto K, Bernhardt N, Brassac J, Mascher M, Hensel G et al. 2019. Unleashing floret fertility in wheat through the mutation of a homeobox gene. Proceedings of the National Academy of Sciences of the United States of America 116: 5182–5187.

Sanchez-Martin J, Steuernagel B, Ghosh S, Herren G, Hurni S, Adamski N, Vrána J, Kubaláková M, Krattinger SG, Wicker T et al. 2016. Rapid gene isolation in barley and wheat by mutant chromosome sequencing. Genome Biology 17: 221.

Shen X, Song S, Li C, Zhang J. 2022. Synonymous mutations in representative yeast genes are mostly strongly non-neutral. Nature 606:725–731.

Sosnowski O, Charcosset A, Joets J. 2012. BioMercator V3: an upgrade of genetic map compilation and quantitative trait loci meta-analysis algorithms. Bioinformatics 28: 2082–2083.

Stanke M, Waack S. 2003. Gene prediction with a hidden Markov model and a new intron submodel. Bioinformatics 19ƒ29(suppl. 2): ii215–ii225.

Steuernagel B, Periyannan SK, Hernández-Pinzón I, Witek K, Rouse MN, Yu G, Hatta A, Ayliffe M, Bariana H, Jones JDG et al. 2016. Rapid cloning of disease-resistance genes in plants using mutagenesis and sequence capture. Nature Biotechnology 34: 652–655.

Su Z, Bernardo A, Tian B, Chen H, Wang S, Ma H, Cai S, Liu D, Zhang D, Li T et al. 2019. A deletion mutation in TaHRC confers Fhb1 resistance to Fusarium head blight in wheat. Nature Genetics 51: 1099–1105.

Sun Z, Zhang M, An Y, Han X, Guo B, Lv G, Zhao Y, Guo Y, Li S. 2022. CRISPR/Cas9-mediated disruption of Xylanase inhibitor protein (XIP) gene improved the dough quality of common wheat. Frontiers in Plant Science 13, 811668 (2022).

Thind AK, Wicker T, Šimková H, Fossati D, Moullet O, Brabant C, Vrána J, Doležel J, Krattinger SG. 2017. Rapid cloning of genes in hexaploid wheat using cultivar-specific long-range chromosome assembly. Nature Biotechnology 35: 793–796.

Uauy C, Distelfeld A, Fahima T, Blechl A, Dubcovsky J. 2006. A NAC gene regulating senescence improves grain protein zinc and iron content in wheat. Science 314: 1298–1301.

Van Ooijen JW. 2009. MapQTL® 6: Software for the mapping of quantitative trait loci in experimental populations of diploid species. (Kyazma BV, Wageningen, Netherlands).

Vercruysse J, Bel MV, Osuna-Cruz CM, Kulkarni SR, Storme V, Nelissen H, Gonzalez N, Inzé D, Vandepoele K. 2019. Comparative transcriptomics enables the identification of functional orthologous genes involved in early leaf growth. Plant Biotechnology Journal18: 553–567.

Wang D, Yu K, Jin D, Sun L, Chu J, Wu W, Xin P, Gregová E, Xin Li X, Sun J et al. 2020. Natural variations in the promoter of Awn Length Inhibitor 1 (ALI-1) are associated with awn elongation and grain length in common wheat. Plant Journal 101: 1075–1090.

Wang H, Zhang Z, Li H, Zhao X, Liu X, Ortiz M, Lin C, and Liu B. 2013. CONSTANS-LIKE 7 regulates branching and shade avoidance response in Arabidopsis. Journal of Experimental Botany 64: 1017–1024.

Wu Y, Bhat PR, Close TJ, Lonardi S. 2008. Efficient and accurate construction of genetic linkage maps from the minimum spanning tree of a graph. PLoS Genetics 4: e1000212.

Xu Y, La G, Fatima N, Liu Z, Zhang L, Zhao L, Chen Mi, Bai G 2021. Precise mapping of QTL for Hessian fly resistance in the hard winter wheat cultivar ‘Overland’. Theoretical and Applied Genetics 134: 3951–3962.

Yan L, Loukoianov A, ranquilli G, Helguera M, Fahima T, Dubcovsky J. 2003. Positional cloning of the wheat vernalization gene VRN1. Proceedings of the National Academy of Sciences of the United States of America 100: 6263–6268.

Yao W, Li G, Yu Y, Ouyang Y. 2018. funRiceGenes dataset for comprehensive understanding and application of rice functional genes. Gigascience 7: 1–9.

Zhang ZL, Ji R, Li H, Zhao T, Liu J, Lin C, Liu B. 2014. CONSTANS-LIKE 7 (COL7) is involved in phytochrome B phyB)-mediated light-quality regulation of auxin homeostasis. Molecular Plant 7: 1429–1440.

Zhang JP, Yu Y, Feng YZ, Zhou YF, Zhang F, Yang YW, Lei MQ, Zhang YC, Chen YQ. 2017. MiR408 regulates grain yield and photosynthesis via a phytocyanin protein. Plant Physiology 175: 1175–1185.

Zhang S, Zhang R, Song G, Gao J, Li W, Han X, Chen M, Li Y, Li G. 2018. Targeted mutagenesis using the Agrobacterium tumefaciens-mediated CRISPR-Cas9 system in common wheat. BMC Plant Biology 18: 1–12.

Zhang X, Jia H, Li T, Wu J, Nagarajan R, Lei L, Powers C, Kan CC, Hua W, Liu Z et al. 2022. TaCol-B5 modifies spike architecture and enhances grain yield in wheat. Science 376: 180–183.

